# Nonmuscle myosin II shRNA inhibit migration and contraction in rat hepatic stellate cells through regulating AKT/mTOR/S6K/4EBP1 signaling pathway

**DOI:** 10.1101/313601

**Authors:** Zhenghong Li, Yun Feng, Ruling Zhang, Peiwen Wang, Lungen Lu, Yuwei Dong

## Abstract

Migration and contraction of activated hepatic stellate cell (HSC) are essential factors for cirrhosis formation and development. It has been demonstrated that blebbistatin, a nonmuscle myosin II (NMMII) inhibitor, can inhibit the migration and contraction of HSC, whereas the main cell signaling pathway is still unknown. Mammalian target of rapamycin (mTOR) signaling pathway may be involved in many cells migration and contraction, whether NMMII and mTOR have any crosslinks draw our attention. In the currently study, we used LV-RNAi to specifically attenuate mTOR and NMMII in rat HSC. We aimed to examine the effect of mTOR LV-RNAi on the migration and contraction of HSC and explore the crosslink between mTOR cell signal and NMMII. Using real-time PCR and western blot, we found that mTOR and the downstream factors including S6K and 4EBP1 all up-regulated with the activation of HSC, mTOR and NMMII LV-RNAi was transfected into activated HSC using lipofectamine 2000. The levels of mRNA and proteins were also examined using real-time PCR and western blot respectively. The expression of mTOR can be down-regulated by NMMII LV-RNAi significantly, as well as the expression of S6K, 4EBP1, α-SMA and collagen I, but the level of AKT was up-regulated. Then we used Transwell system and collagen lattices to examine the NMMII and mTOR LV-RNAi efficiency on HSC migration and contraction, as we hypothesized, both of the LV-RNAi could inhibit HSC migration and contraction significantly. These results indicated that nonmuscle myosin II shRNA inhibit migration and contraction in rat hepatic stellate cells through the regulation of mTOR/S6K/4EBP1 signaling pathway

## 1. Introduction

Chronic injury leading to liver fibrosis occurs in response to a variety of insults, including viral hepatitis, alcohol abuse, drugs, metabolic diseases due to overload of iron or copper, autoimmune attack of hepatocytes or bile duct epithelium, or congenital abnormalities^[1]^.Typically, injury is existed for months to years before significant scar accumulates, although the time course may be accelerated in congenital liver disease. Liver fibrosis is reversible, whereas cirrhosis, the end-stage consequence of fibrosis, is generally irreversible^[2]^.

Under conditions of liver injury, hepatic stellate cells (HSC) undergo a change in phenotype, called activation. These phenotypic changes, are essential for both wound healing and fibrosis of the liver^[3]^. The pathophysiologic importance of HSC migration is supported by the observation that HSC migrate to and accumulate in areas of injury and fibrosis remote from their usual location encircling sinusoids during wound healing and cirrhosis^[4, 5]^.

In addition to altering the wound healing, HSC hypercontractility contributes to increased resistance of sinusoids manifesting portal hypertension, characterized by both increased portal blood flow and intrahepatic vascular tone^[1, 6, 7]^. Portal hypertension is the result of augmented intrahepatic vascular resistance and increased portal blood flow. Accumulating evidences from in vitro and in vivo studies suggest that stellate cells are also involved in the regulation of the liver microcirculation and portal hypertension. Activated hepatic stellate cells have the necessary machinery to contract or relax in response to a number of vasoactive substances^[8]^.

HSC migration and contraction are necessary for the wound-healing process and influence both development and severity of hepatic fibrosis. Recent studies examined the expression and functionally of nonmuscle myosin II (NMMII) protein in mouse HSCs^[9]^. Inhibition of myosin II ATPase by blebbistatin, a cell-permeable pharmacological agent, altered HSC morphology and reduced characteristic HSC contraction. However, the cell signal of how NMMII inhibit the migration and contraction of HSC has been unknown.

Several evidence indicated that the phosphatidylinositol 3-kinase (PI3K)/the mammalian target of rapamycin (mTOR) signaling pathway may be involved in many cells migration and contraction mTOR activates ribosomal S6 kinase (S6K1 and S6K2) and eukaryotic initiation 4E (4EBP1) to regulate cell-cycle progression and protein synthesis^[10]^. Whereas, whether mTOR cell signal also influence the migration and contraction of HSC has been unknown, and whether NMMII and mTOR have any interaction has to be investigated. The aim of this study was to evaluate the effects of mTOR in HSC migration and contraction, more importantly, we want to find the relationship between NMMII and mTOR, therefore, we can clarify the cell signal of NMMII in regulating the migration and contraction of HSC.

## 2. Materials and methods

### 2.1 Cell isolation and identify

Primary HSCs were isolated from livers of Wistar rats and cultured on plastic dishes in Dulbecco’s modified Eagle’s medium (DMEM;Invitrogen), supplemented with 4 mmol/L L-glutamine, 10% FCS, and penicillin (100IU/ml)/streptomycin (100mg/ml). The viability of the isolated cells was determined using trypan blue staining. The purity of isolated quiescent HSCs was determined by vitamin A autofluorescence and routinely exceeded 90%. HSC-T6 was also enrolled as control.

### 2.2 Real-time PCR

Total RNA was extracted from cells using Trizol and was reverse-transcribed using an iScript cDNA kit (Takara). Real-time PCR was performed on an iCycler system using SYBR Green Master Mix (Takara). Primer specificity was confirmed by sequencing PCR products. β-actin was used as the internal control. Data were presented according to the ΔΔC_T_ method.

### 2.3 Western blot analysis

Cultured cells were homogenized in ice-cold RNA buffer containing protease and phosphatase inhibitors (Sigma). Cell lysate was separated on 10% Bis-Tris gradient gel (Invitrogen), transferred to a PVDF membrane, then incubated with antibody to NMMII, mTOR, S6K1, S6K2, 4EBP1, α-SMA, protein kinase B (AKT), COLI, COLIII (Abcam). GAPDH was used as a loading control.

### 2.4 Contraction assay

Collagen lattices were prepared and culture-activated HSCs were seeded onto the congealed collagen lattice and allowed to recover overnight. Collagen lattices were dislodged from wells with a 10μL pipette tip. The differences in collagen diameters were reported as percentage change in collagen lattice circumference.

### 2.5 Migration assay

Cell motility was determined in vitro firstly using a Transwell chamber (BD). Cells were trypsinized and placed into the upper wells of the Boyden chamber (10000 cells per ml) in 100μ DMEM with 20% FBS. In the lower chamber, 600μl DMEM containing 10% FBS was added. Cells in the Boyden chamber were incubated for 48 hours at 37°C in a 5% CO2 incubator. After non-migrated cells were scraped off, while the cells in the bottom well were collected and counted by crystal violet stain.

### 2.5 Lentiviral vectors for NMMII and mTOR shRNA

Small hairpin RNA (shRNA) targeting rat NMMII and mTOR were designed as follows. The sense of NMMII was: 5’- CACCGCCTCCACAAGACATGCGTATTCGAAAATACGCATGTCTTGTGGATT-3’ the antisense was: 5’- AAAACCTCCACAAGACATGCGTATTTTCGAATACGCATGTCTTGTFFAGGC -3’. The sense of mTOR was: 5’- CACCGGTCATGGTCATGCCCACGTTCCTTAACGAATTAAGGAACGTFFFCA TGACC-3’ the antisense was: 5’- CACCGGTCATGCCCACGTTCCTTAACGAATTAAGGAACGTGGGAACGTGGGCATGACC-3’.

The recombinant lentivirus gene transfer vector targeting NMMII and mTOR pGCSIL-GFP-NMMII/mTOR (LV-RNAi) encoding the green fluorescent protein (GFP) sequence was constructed as previously described. The targeting sequence of the shRNA was confirmed by sequencing. The lentiviral vector pGCSIL-GFP-Negative (LV-NC) containing an invalid RNAi sequence was used to monitor non-specific responses caused by heterologous siRNA. The LV-RNAi and the LV-NC were prepared to 5*10^9^ Tu/ml.

### 2.6 Lentiviral vector transfection

Cells were subcultured at 5*10^4^ cells/well into 6-well tissue culture plates overnight. The viral supernatant was then added into cells at a multiplicity of infection (MOI) of 10 with 10% FBS and 8μg/ml polybrene. The infected cells were considered to be the LV-RNAi and the LV-NC group, respectively, and the cells without infection were considered as the control group. Flow cytometry was used to detect the transfection efficiency, and fluorescence microscopy was used to observe the cells which release fluorescence.

### 2.7 Statistical analysis

Data are the mean ± SD, and are representative of at least 3 separate experiments. Comparisons were performed using ANOVA, and correlations were determined using Pearson’s correlation coefficient. A P-value of less than 0.05 was considered statistically significant.

## 3. Results

### 3.1 Differential expression of NMMII and mTOR in quiescent and activated HSCs

Primary rat HSCs were isolated and culutued for 0, 24h and 72h. The mRNA and protein levels of NMMII, mTOR, AKT, 4EBP1, S6K1 and S6K2 were evaluated using real-time PCR and western blot assays. We observed that the expression of NMMII, mTOR, AKT, 4EBP1, S6K1 and S6K2 all gradually increased 1.5-4 fold compared with quiescent in both mRNA and protein level (Fig1).

**Figure 1.**
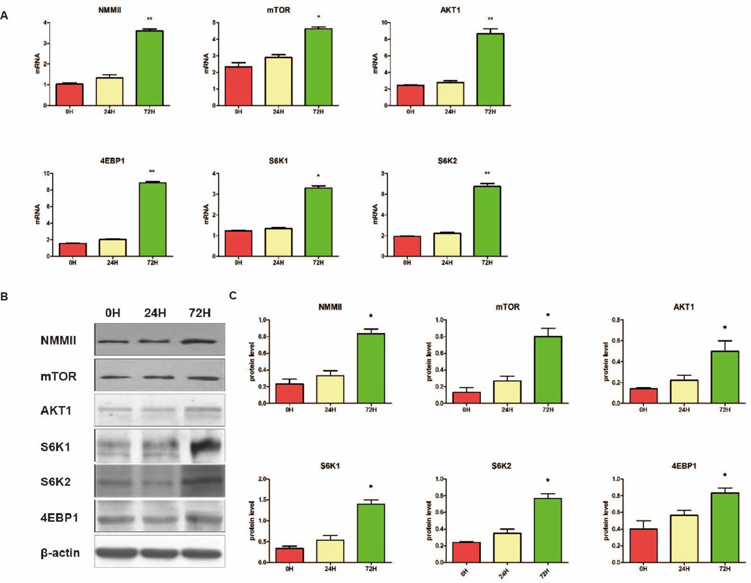
NMMII and mTOR were upregualted in activated HSCs. A: mRNA expression of NMMII, mTOR, AKT1, S6K1, S6K2 and 4EBP1 were assessed in quiescent and activated HSCs (0H, 24H and 72H) by real time PCR. mRNA expression of the above genes were normalized to total cDNA concentration.. B and C: Protein expression of NMMII, mTOR, AKT1, S6K1, S6K2 and 4EBP1 in quiescent and activated HSCs (0H, 24H and 72H) by Western blot analysis. (* *p*<0.05 as compared to quiescent).

### 3.2 The interaction of NMMII and mTOR in activated HSCs

The effects of NMMII and mTOR shRNA on the mRNA and protein level of NMMII and mTOR were determined by real-time PCR and western blot assays. In NMMII LV-RNAi group, the total mRNA of NMMII, mTOR, α-SMA, S6K1, S6K2, 4EBP1, COLI and COLIII decreased to 22%, 50%, 60%, 55%, 60%, 40%, 60% and 55% respectively, after 48 hours transfection. While the mRNA level of AKT was up-regulated to 1.5 fold (Fig 2A). Western blot assays revealed that the protein level of NMMII, mTOR, α-SMA, S6K1, S6K2, 4EBP1, COLI and COLIII also down-regulated by NMMI LV-RNAi and had statistical significance(Fig 2B-C)

**Figure 2.**
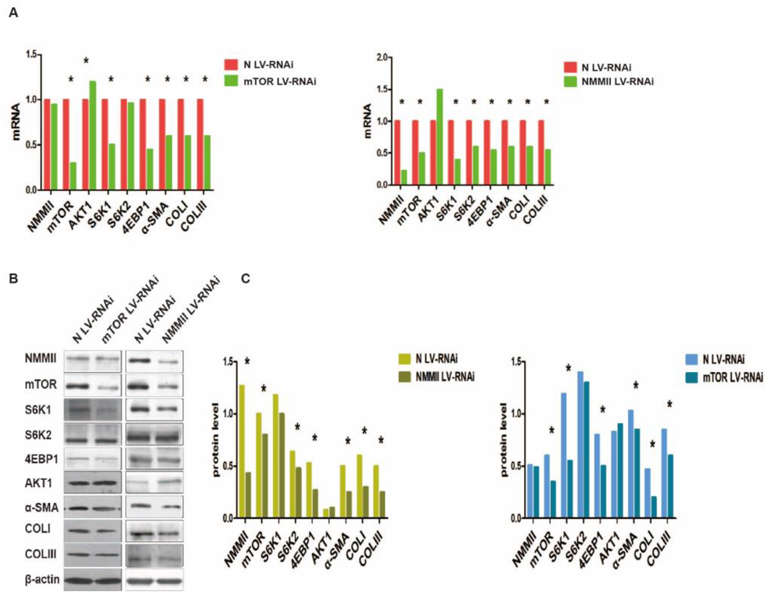
NMMII and mTOR interaction and NMMII, mTOR LV-RNAi influence HSC secretion ECM. Activated HSCs were transfected with NMMII and mTOR LV-RNAi or N LV-RNAi as control. A: Real Time PCR determined gene-specific inhibition of each genes as normalized to total cDNA concentration compared to scramble control. B and C: The NMMII, mTOR, S6K1, S6K2, 4EBP1, AKT1, α-SMA, COLI and COLIII protein level of HSCs treated with NMMII and mTOR LV-RNAi were examined by WB. The values represent ration of β-actin. (* p<0.05 as compared to quiescent).

In the mTOR LV-RNAi group, the total mRNA of mTOR, α-SMA, S6K1, 4EBP1, COLI and COLIII decreased to 30%, 60%, 50%, 45%, 60% and 60% respectively. While the mRNA level of NMMII and S6K2 had no significant difference, but the AKT mRNA level was also up-regulated to 1.2 fold (Fig 2A). Western blot assay results were accordance with that of real-time PCR (Fig 2B-C).

### 3.3 Effect of mTOR LV-RNAi and rapamycin on HSCs migration and contraction

To determine functional contributions of mTOR in HSC migration, culture-activated cells (Day 3) were treated with mTOR LV-RNAi and rapamycin (0.5nmol/L) for 24 hours. The effect of mTOR LV-RNAi and rapamycin were examined using the Transwell assay. We found that the treatment of mTOR LV-RNAi and rapamycin both can inhibit the migration of activated HSCs significantly (Fig 3A).

To investigate the function of mTOR in HSCs contration, HSCs were treated with mTOR LV-RNAi and rapamycin (0.5 nmol/L) and test the changes of gel circumference after 24h as compared to scramble control. Results indicated that LV-RNAi and rapamycin both can inhibit HSCs contraction significantly (Fig 3B).

**Figure 3.**
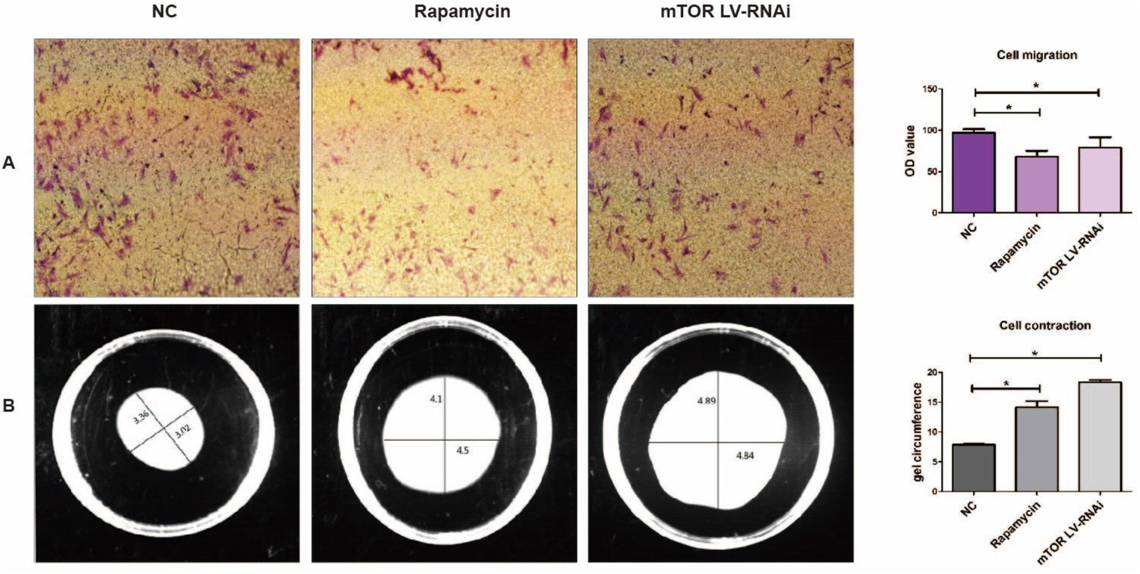
Effect of mTOR LV-RNAi on HSC migration and contraction. Activated HSCs were transfected with LV-RNAi targeted to mTOR or NC LV-RNAi as control for 48h. A: Migration activaties were measured using the Transwell system. Cells that migrated were stained with 0.05% Crystal violet. B:HSCs contraction was quantified using PTI ImageMaster software and reported as percentage change in gel cirumference. (* p<0.05 as compared to control).

### 3.4 Effect of NMMII LV-RNAi and blebbistatin on HSCs migration and contraction

The function of NMMII on HSC migration was also tested with Transwell assay. HSCs (Day 3) were transfected with NMMII LV-RNAi and blebbistatin (10umol/L) for 24 hours separately. The OD values revealed that treatment of NMMII LV-RNAi and blebbisatin can significantly inhibit the migration of HSCs. To find the changes of HSCs treated with NMMII LV-RNAi or blebbistatin, collagen lattices was enrolled. After 24 hours treated with NMMII LV-RNAi or blebbistatin (10umol/L), the gel circumference were examined to reveal the differences. The outcomes told us that NMMII LV-RNAi and blebbistatin both can inhibit the construction of HSCs.(Fig4)

**Figure 4.**
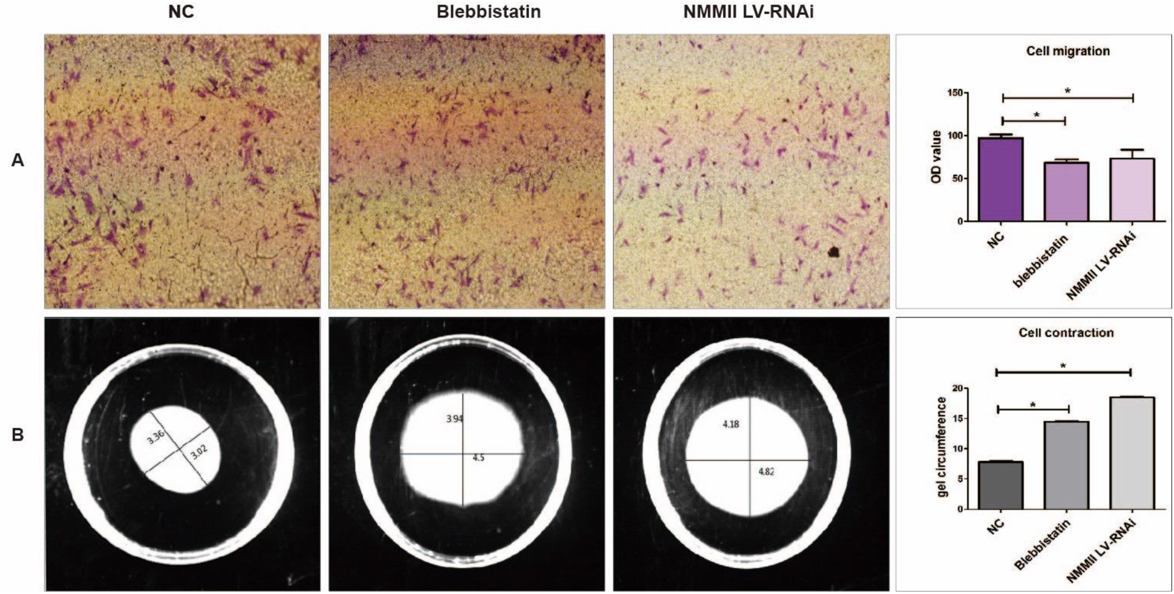
Effect of NMMII LV-RNAi on HSC migration and contraction. Activated HSCs were transfected with LV-RNAi targeted to NMMII or NC LV-RNAi as control for 48h. A: Migration activaties were measured using the Transwell system. Cells that migrated were stained with 0.05% Crystal violet. B:HSCs contraction was quantified using PTI ImageMaster software and reported as percentage change in gel cirumference. (* p<0.05 as compared to control).

## 4. Discussion

The main findings in this study are: 1) mTOR/S6K/4EBP1 cell signaling were activated along with the activation of HSC; 2) mTOR LV-RNAi could inhibit migration and contraction in activated HSC; 3) mTOR/S6K/4EBP1 cell signal were down-regulated by NMMII LV-RNAi. Therefore, we came to the conclusion that NMMII LV-RNAi can inhibit migration and contraction in rat hepatic stellate cells through regulating AKT/mTOR/4EBP1/S6K cell signaling pathway.

HSC are located in the perisinusodial space of Disse between the hepatocyte and endothelial cells^[11]^. In diseased liver, chemotactic factors released during injury stimulate HSC migration to damaged areas. If the damage persistence, hyper contraction of HSC will lead to increased portal blood flow and intrahepatic vascular tone^[12]^. mTOR/S6K/4EBP1 cell signal has been certified that this cell signal can inhibit several cell types’ migration and contraction, such as HL-60, HLE B3, et al^[10, 13]^. Based on these findings, we test the expression differences of mTOR cell signal in the process of HSC activation. According to previous findings, mTOR, S6K1, S6K2 and 4EBP1 were all up-regulated with the activation of HSC.

mTOR is the core component of two distinct complexes only partially characterized: complex 1 (mTORC1) and complex 2 (mTORC2). mTOR can be specifically inhibited by rapamycin only when it is in mTORC1, leading to the initial definition of mTORC1 as “rapamycin sensitive” and of mTORC2 as “rapamycin insensitive”^[14]^. In our experiment, activated HSC treated with 0.5 nmol/L rapamycin, caused the migration and contraction ability of HSC decreased significantly. According to these results, we can conclude that in HSC, mTOR was targeted with mTORC1.

The two effectors of mTORC1 that are better characterized are the ribosomal S6 kinases (S6K1 and S6K2) and the eukaryotic initiation factor 4 binding proteins 1(4E-BP1). S6K is activated by phosphorylation at several sites. Upon activation, S6K phosphorylates subunit 6 of ribosomal protein, leading to general activation of translation through an unclear and still highly debated mechanism. Phosphorylation of 4EBP instead leads to dissociation from eukaryotic initiation factor 4E, which is released, allowing the formation of translation initiation complexes^[15]^.

In this article, we enrolled LV-RNAi technique to clarify the relationship of mTOR with S6K/4EBP1 and NMMII. Firstly, our results revealed that mTOR LV-RNAi can down-regulated the expression of S6K1 and 4EBP1 but had no influence on S6K2 and NMMII in activated HSC. S6K and 4EBP1 were the downstream factors of mTOR, therefore, the expression change of mTOR can influence the expression of S6K1 and 4EBP1. However, the S6K2 level had no difference indicated that S6K2 was not as sensitive as S6K1 to the change of mTOR expression. More importantly, NMMII level was not changed with the different expression of mTOR, this consequence indicated that NMMII was not controlled by mTOR, thus the cell signal of NMMII has to clarify.

In order to identify the cell signal of NMMII to regulate the migration and contraction of activated HSC, we used NMMII LV-RNAi and blebbistatin (10umol/L) to infect activated HSC, we found NMMII LV-RNAi and blebbistatin both can significant inhibit the migration and contraction of HSC, the influence of the two groups had no difference. Then, we also enrolled NMMII LV-RNAi technique to identify the alteration of NMMII on the expression of mTOR and downstream factors. The results of this experiment indicated that NMMII LV-RNAi could down-regulate the expression of mTOR, S6K1, S6K2 and 4EBP1, but up-regulated the expression of AKT. According to this result, we can conclude that NMMII can influence the migration and contraction of HSC through mTOR/S6K/4EBP1 cell signaling pathway, and as an important upstream factor, the level of AKT was up-regulated. It has been described that AKT is the central molecule in the AKT/mTOR pathway, activating and modulating numerous downstream targets. AKT can stimulate protein synthesis by activating mTOR through inhibition of the Tuberous sclerosis complex (TSC1/2)^[10]^. In addition, according to our research, not only mTOR but also NMMII could be regualted by AKT. Because in both NMMII and mTOR LV-RNAi group, the expression of AKT were upregualted. Although the upregualtion is not significant, we still believe that the upregualtion of AKT was a degenerative feedback by NMMII and mTOR downregulated. In addition, we had also found that NMMII and mTOR LV-RNAi also can down-regulated the expression of α-SMA and COLI. HSC are located in the perisinusodial space of Disse between the hepatocyte and endothelial cells^[2]^. In diseased liver, damaging stimuli trigger quiescent HSC to an activated myofibroblast-like cell. Activated HSC increase expression of cytoskeletal protein such as α-SMA and secrete numerous ECM protein including type I collagen leading to disruption of normal liver architecture impeding liver microcirculation. The cytoskeletal and ECM protein changes maybe the key factors leading to HSC get the migration and contraction ability^[16]^.

In conclusion, our study clearly demonstrated that as NMMII can regulate the expression of α-SMA and COLI through mTOR/S6K/4EBP1 cell signal, NMMII LV-RNAi can inhibit the migration and contraction of HSC.

## Ackonwledements

This study was supported by the National Natural Science Foundation of China (81070345 & 81300326) and Chinese foundation for hepatitis prevention and control Tian Qing liver disease foundation (TQGB2011024)

